# Integrated Diffusion Image Operator (iDIO): A tool for automated configuration and processing of diffusion MRI data

**DOI:** 10.1101/2022.03.14.483870

**Authors:** Chih-Chin Heather Hsu, Shin Tai Chong, Yi-Chia Kung, Kuan-Tsen Kuo, Chu-Chung Huang, Ching-Po Lin

**Affiliations:** Institute of Neuroscience, National Yang Ming Chiao Tung University, Taipei, 112 Taiwan; Center for Geriatrics and Gerontology, Taipei Veterans General Hospital, Taipei, 112 Taiwan; Shanghai Key Laboratory of Brain Functional Genomics (Ministry of Education), Affiliated Mental Health Center (ECNU), School of Psychology and Cognitive Science, East China Normal University, Shanghai 200062, China; Shanghai Changning Mental Health Center, Shanghai 200335, China; Brain Research Center, National Yang Ming Chiao Tung University, Taipei, 112 Taiwan

**Keywords:** Preprocessing, pipeline tool, diffusion MRI, connectome

## Abstract

The preprocessing of diffusion magnetic resonance imaging (dMRI) data involves numerous steps, including the corrections for head motion, susceptibility distortion, low signal-to-noise ratio, and signal drifting. Researchers or clinical practitioners often need to configure different preprocessing steps depending on disparate image acquisition schemes, which increases the technical threshold for dMRI analysis for non-expert users. This could cause disparities in data processing approaches and thus hinder the comparability between studies. To make the dMRI data processing steps transparent and adapt to various dMRI acquisition schemes for researchers, we propose a semi-automated pipeline tool for dMRI named integrated Diffusion Image Operator or iDIO. This pipeline integrates features from a wide range of advanced dMRI software tools and targets at providing a one-click solution for dMRI data analysis, via automatic configuration for a set of optimal processing steps based on the image header of the input data. Additionally, the pipeline provides options for post-processing, such as estimation of diffusion tensor metrics and whole-brain tractography-based connectomes reconstruction using common brain atlases. The iDIO pipeline also outputs an easy-to-interpret quality control report to facilitate users to assess the data quality. To keep the transparency of data processing, the execution log and all the intermediate images produced in the iDIO’s workflow are accessible. The goal of iDIO is to reduce the barriers for clinical or non-specialist users to adopt the state-of-art dMRI processing steps.

## Introduction

Diffusion magnetic resonance imaging (dMRI) is the primary noninvasive in vivo technique for investigations of white matter fiber properties and the structural network of the human brain, such as the voxel-wise tract-based spatial statistics using diffusion tensor metrics (e.g. fractional anisotropy and mean diffusivity) (Smith et al., 2006), or measuring the topological properties of the complex human brain organization using tractography-based analysis (Bastiani and Roebroeck, 2015; Yeh et al., 2021; Zhang et al., 2021). Since the last decade, dMRI technologies have been rapidly advanced in both hardware and software. Clinical dMRI scans limited by the scanner capability and scanning time may involve restrictions on imaging protocols and parameters, a guideline or recommendation is urgently needed to make dMRI data preprocessing more standardized (Veraart and Descoteaux, 2021).

For diffusion-weighted images (DWIs) acquired using full k-space coverage, multiple non-diffusion-weighted (i.e. b=0) images distributed equally, and without image interpolation, applying data preprocessing such as signal denoising, Gibbs artifact removing, and signal drifting correction may improve the precision of diffusion-related estimates (Kellner et al., 2016; Veraart et al., 2016a; Veraart et al., 2016c; Vos et al., 2017). Ideally, these steps shall be taken prior to common dMRI preprocessing including brain extraction, susceptibility distortion correction, eddy current, and motion correction (Smith, 2002; Andersson et al., 2003; Andersson and Sotiropoulos, 2016). On the other hand, when the dMRI acquisition is composed of multiple b-values or so-called multi-shells, it would be convenient if a processing pipeline can automatically determine the suitable diffusion models, for instance, extracting low b-value shell for the diffusion tensor analysis while using the whole multi-shell data for the reconstruction of fiber orientation distributions (Tournier et al., 2013).

Advanced software packages have provided individual functions to conduct specific or combined data processing, such as ANTs, FSL, PreQual, and MRtrix3 (Avants et al., 2009; Jenkinson et al., 2012; Tournier et al., 2019; Cai et al., 2021). Several processing pipelines have also been proposed to deal with the variations in image acquisition schemes (Smith and Connelly, 2019; Theaud et al., 2020; Cai et al., 2021; Cieslak et al., 2021; Joseph et al., 2021). These tools can improve the comparability or generalizability of the research findings across studies. However, the preprocessing steps sometimes require sophisticated user settings or restricted execution order. Moreover, the quality control (QC) for the preprocessed images is usually done either separately or only for specific steps (Bastiani et al., 2019). Hence, QC is often omitted or unaware by the end-users. Lastly, a semi-automated pipeline that can deliver a range of dMRI-based measures may facilitate an end-user’s downstream applications, such as those used in the voxel-wise analysis, fixel-wise analysis, white matter tract profiling or so-called tractometry, and brain network analysis (Smith et al., 2006; Behrens et al., 2007; Raffelt et al., 2017b).

An adaptive pipeline that integrates the current state-of-the-art dMRI data preprocessing methods and delivers essential QC reports may benefit the neuroimaging community via lowering the technical requirements for researchers. Here, we present the ‘integrated Diffusion Imaging Operators’ or iDIO, which aims to be a user-friendly one-click solution for dMRI data processing based upon advanced software tools. The workflow of iDIO encapsulates parts of the functionalities of ANTs (Avants et al., 2009), FSL (Jenkinson et al., 2012), MRtrix3 (Tournier et al., 2019), and Prequal (Cai et al., 2021). iDIO supports dMRI data acquired with different clinical MRI vendors, such as Siemens, GE, Philips, and United Imaging. Based on the DICOM header of the input data, iDIO automatically generates a configuration file that determines the appropriate preprocessing steps. All the steps are transparent in that users can inspect all the input, interim, and output images, alongside a log file that lists the steps taken in a run. iDIO uses a setting file that includes all options and parameters information, to minimize using erroneous input options and provide better consistencies across studies. iDIO includes functionalities to deliver basic dMRI measures and structural connectivity matrices with multiple brain atlases.

## Methods

### iDIO workflow

iDIO is programmed with bash shell and python scripts to conduct image preprocessing and analysis. Figure 1 shows the summary of iDIO’s workflow. iDIO accepts data provided in the BIDS format (Gorgolewski et al., 2016). The configuration of processing steps majorly relies on the image JSON file to reduce manual error during parameters configuration. iDIO collects image acquisition information by reading the JSON tags and then automatically adapts the preprocessing steps to be done. A QC report will be provided and preprocessed images were kept in each step. In the following sections, we sequentially described the details of each processing step.

**Figure 1.**
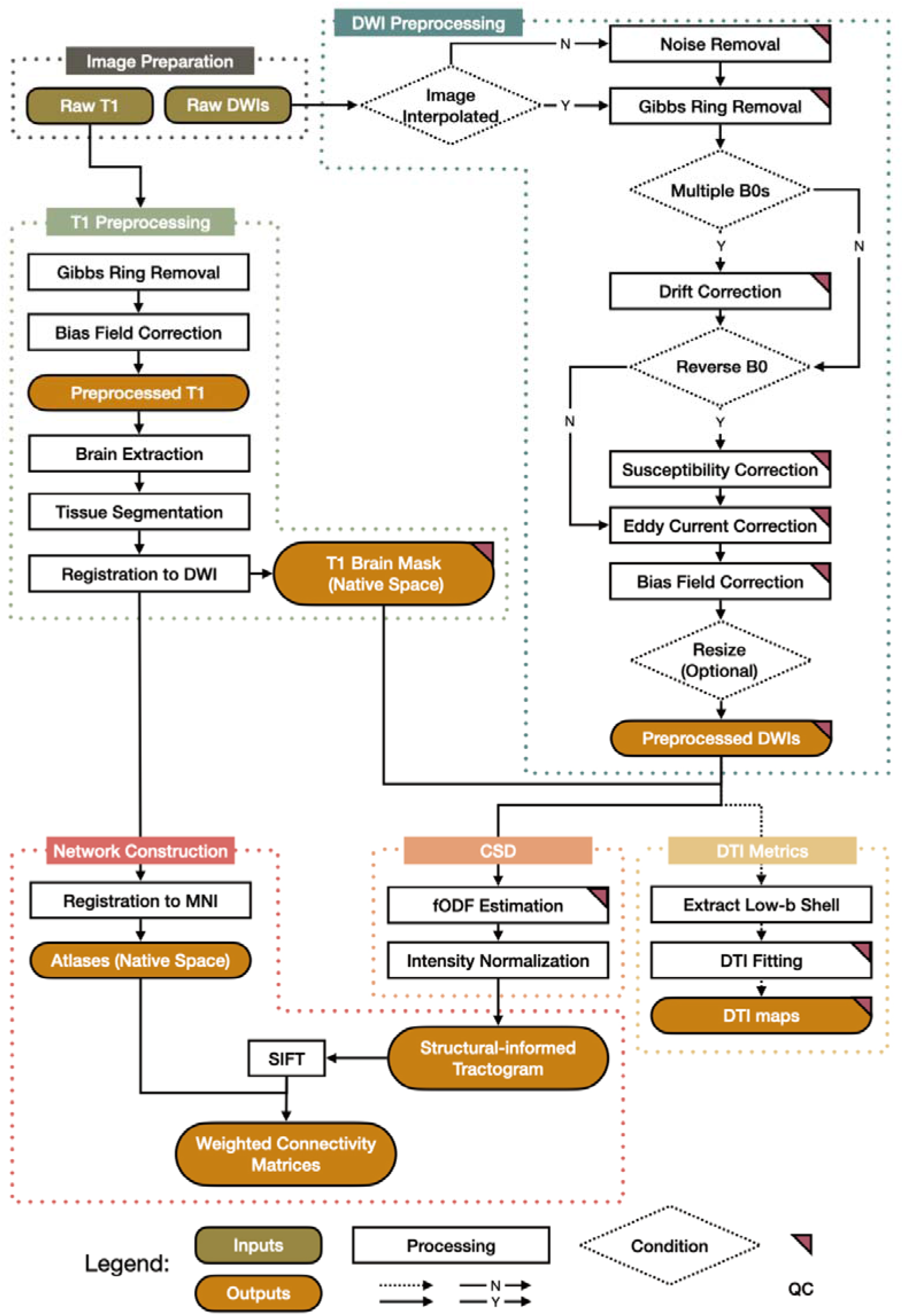
The flowchart of the main process of the iDIO pipeline. iDIO is composed of six main blocks (image preparation, DWI preprocessing, T1 Preprocessing, DTI Metrics, CSD and network construction). Images are preprocessed according to the black arrow direction. T1 image is essential for generating brain masks and providing anatomical reference for diffusion analyses. DTI: diffusion tensor image; CSD: constrained spherical deconvolution; SIFT: spherical-deconvolution informed filtering.

#### 1 DWIprep: renaming, and generating description file

First, iDIO checks for the (in)consistency between the file name and the phase encoding direction of the DWIs; it will rename the input DWIs with phase encoding information. Next, iDIO will generate four configuration files for the subsequent processing steps; they are 1) phase encoding information (Index_PE.txt); 2) image acquisition parameter (Acquparams_Topup.txt); 3) an index file (Eddy_index.txt) that indicates the relevant data corresponding to Acquparams_Topup.txt file, and a file (MBF.txt) contains the multi-band factor (Andersson et al., 2003; Smith et al., 2004; Jenkinson et al., 2012).

#### 2 BiasCo: Noise removal, Gibbs ring removal, and signal drift correction

The preprocessing begins with image denoising (MRtrix3 command: dwidenoise), which includes a noise level estimation and a Marchenko-Pastur principal component analysis (MP-PCA) denoising (Veraart et al., 2016b; Veraart et al., 2016c). To use MP-PCA, DWIs should not be treated with any other data processing since that would change the noise characteristics. However, some vendor machines by default interpolate the reconstructed images to higher spatial resolution, which may unfavorably change the noise distribution. Therefore, if image interpolation is detected (based on image header), the user is notified to skip DWI denoising to avoid incorrect estimation of the raw noise distribution. Second, Gibbs ringing removal is performed by the mrdegibbs command in MRtrix3, which removes the Gibbs ringing artifacts by re-interpolating DWIs based on local, subvoxel-shifts (Kellner et al., 2016). Third, if the DWI sampling scheme contains more than three evenly distributed b=0 images, a linear signal drift correction is then performed to reduce the effect of signal dropping caused by the temporal scanner instability, using the method proposed by Vos et al (2017).

#### 3 EddyCo: Correction for susceptibility-induced distortion, eddy current-induced distortion and subject movement; bias correction and resampling

This step first deals with image distortion caused by the off-resonance field induced by susceptibility and eddy currents, and subject movement, using the topup and eddy functions implemented in FSL (Smith et al., 2004). If there are b=0 images with opposite phase-encoding directions, topup is applied (correcting susceptibility-induced distortion), followed by eddy (correcting for eddy-current-induced distortion and motion). Otherwise, topup is skipped. iDIO automatically looks for the availability of the GPU (CUDA) toolkit on the computer system and selects the corresponding version of eddy_cuda. After eddy, a B1 field inhomogeneity correction is performed using MRtrix3’s dwibiascorrect, based on the N4ITK algorithm (Tustison et al., 2010). This step ends with imaging resampling, which depends on users’ applications and therefore is optional.

#### 4 T1preproc: T1w image preprocessing

T1w image preprocessing includes Gibbs-ringing artifact removal (by mrdegibbs) and B1 field inhomogeneity bias correction (by N4BiasFieldCorrection command in ANTs), both of which can improve skull stripping. A skull-stripped brain mask is generated using the antsBrainExtraction in ANTs, and the five-tissue type (5tt) segmentation images are produced using MRtrix3’s 5ttgen (default using FSL’s methods) (Smith, 2002; Smith et al., 2004; Patenaude et al., 2011; Smith et al., 2012). Next, image registration from T1 to DWI uses the boundary-based registration method (FLIRT in FSL) (Jenkinson and Smith, 2001; Jenkinson et al., 2002). T1w brain mask is then transformed into DWI space for further usage. Tissue segmentation in DWI space is conducted to produce tissue partial volume maps required for anatomically-constrained tractography in step 7_NetworkProc.

#### 5 DTIFIT: diffusion tensor fitting

DWIs with b-values lower than 1500 s/mm^2^ are extracted and analyzed using the diffusion tensor model with weighted least square, done by FSL’s command dtifit (Basser et al., 1994). Eigenvalues, eigenvectors and diffusivity measures (fractional anisotropy, mean diffusivity, axial diffusivity, radial diffusivity,) were provided.

#### 6 CSDpreproc: CSD model fitting

The iDIO pipeline uses CSD to generate fiber orientation distributions (FODs) as a basis for tractography. For both single- and multi-shell data, tissue response functions for white matter (WM), grey matter (GM), and cerebrospinal fluid (CSF) were calculated by MRtrix3’s dwi2response function with the dhollander algorithm (Dhollander et al., 2016). FODs were than be estimated by MRtrix3’s dwi2fod function with multi-shell multi-tissue CSD algorithm (Jeurissen et al., 2014); a two-tissue (WM and CSF) and a three-tissue (WM, GM, and CSF) CSD is used for the single-shell and multi-shell DWIs, respectively. All the generated tissue components will further undergo a multi-tissue informed log-domain intensity normalization to make the component is quantitative across subjects using the MRtrix3’s mtnormalise command (Raffelt et al., 2017a; Tournier et al., 2019).

#### 7 NetworkProc: CSD FOD-based tractography and network construction

iDIO utilizes the WM FODs to generate the whole-brain tractogram with MRtrix3’s tckgen command. Ten million (optional) streamlines are generated using the anatomically-constrained tractography (Smith et al., 2012) with dynamic seeding (Smith et al., 2015a). The spherical-deconvolution informed filtering of tractogram (SIFT) is then applied using tcksift2 command (Smith et al., 2015b). Tractogram-based connectomes are generated using a range of atlases, including automatic anatomical labeling atlas 3 (Rolls et al., 2020), HCP-MMP (Glasser et al., 2016), HCP-MMP with subcortical regions (HCPex) (Huang et al., 2021) and Yeo 400 (Yeo et al., 2011). These atlases are registered with ICBM152’s template T1-weighted images using antsRegistrationSyNQuick script in ANTs (Avants et al., 2009; Fonov et al., 2009) and then transformed to individual’s image space. Three weighted connectivity matrices are generated using (1) weighted streamline count (sum of the SIFT2 track weight) (Smith et al., 2015a), (2) weighted streamline count scaled by proportionality coefficient (Smith et al., 2013) and (3) mean streamline length (Figure 2). The SIFT2 algorithm permits streamline connection densities to be more biologically relevant (Smith et al., 2015a; Yeh et al., 2016).

**Figure 2.**
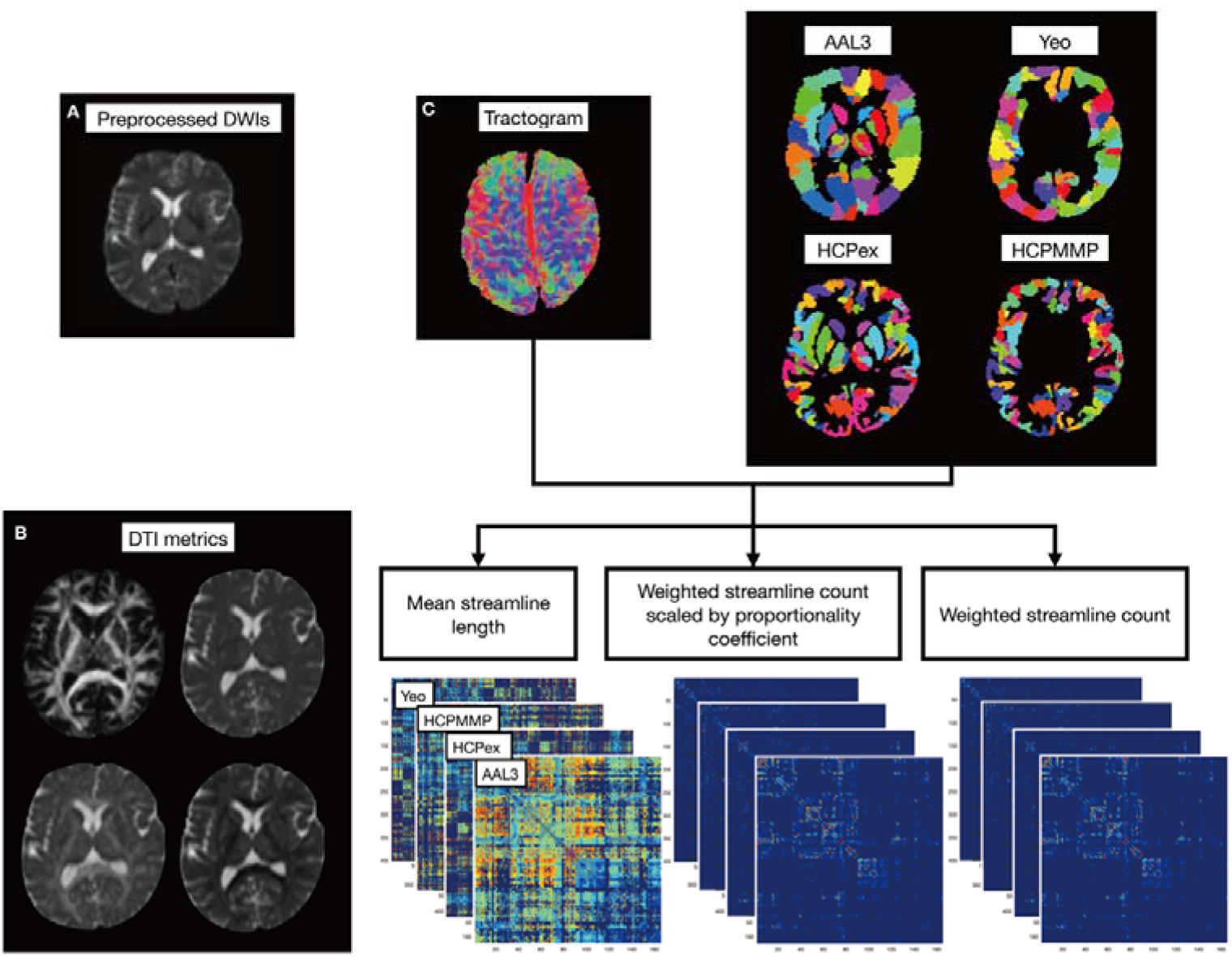
iDIO provides the preprocessed DWIs with DTI metric maps and the brain network using CSD-based tractography. (A) Preprocessed DWIs (B) DTI metrics maps (top left: Fractional anisotropy; top right: mean diffusivity; bottom left: axial diffusivity; bottom right: radial diffusivity) (C) White matter tracks reconstructed using ACT with iFOD2 algorithm. (D) Brain parcellation (nodes) based on pre-defined atlas (AAL3, Yeo, HCPex, HCPMMP) in native space. (E) Generated network matrices weighted by average length, SIFT scaled by mu and SIFT. DTI: diffusion tensor imaging; SIFT: spherical-deconvolution informed filtering of tractogram

### iDIO QC reports

iDIO produces the QC report for each dataset in a single PDF file when preprocessing is completed. Most of the dMRI preprocessing steps have corresponding QC results (Figure 1 with QC labels), which were generated mostly from EddyQuad and PreQual (Bastiani et al., 2019; Cai et al., 2021). The QC report embeds a method summary, and warning messages are given when applicable. Images with severe motion are specifically mentioned in the report (Figure 3A). iDIO QC provides comparisons of DWIs before and after denoising, Gibbs removal, drift correction, b-vector reorientation and bias field correction. The assessments for Eddy/motion correction and SNR/CNR estimation are generated using eddy_quad. CSD FODs and DTI directional encoded color (DEC) maps are given to have a quick check on improperly oriented gradients tables. DTI DEC and the sum of square error (SSE) of tensor fit were generated to help quick visual detections for possible signal dropout and chemical shift artifacts (Figure 3B and 3C). Voxels having higher diffusion-weighted signal than the b=0 intensity are output in an additional image volume. A complementary CSV document is provided for screening the quality of image preprocessing for a group of datasets.

**Figure 3.**
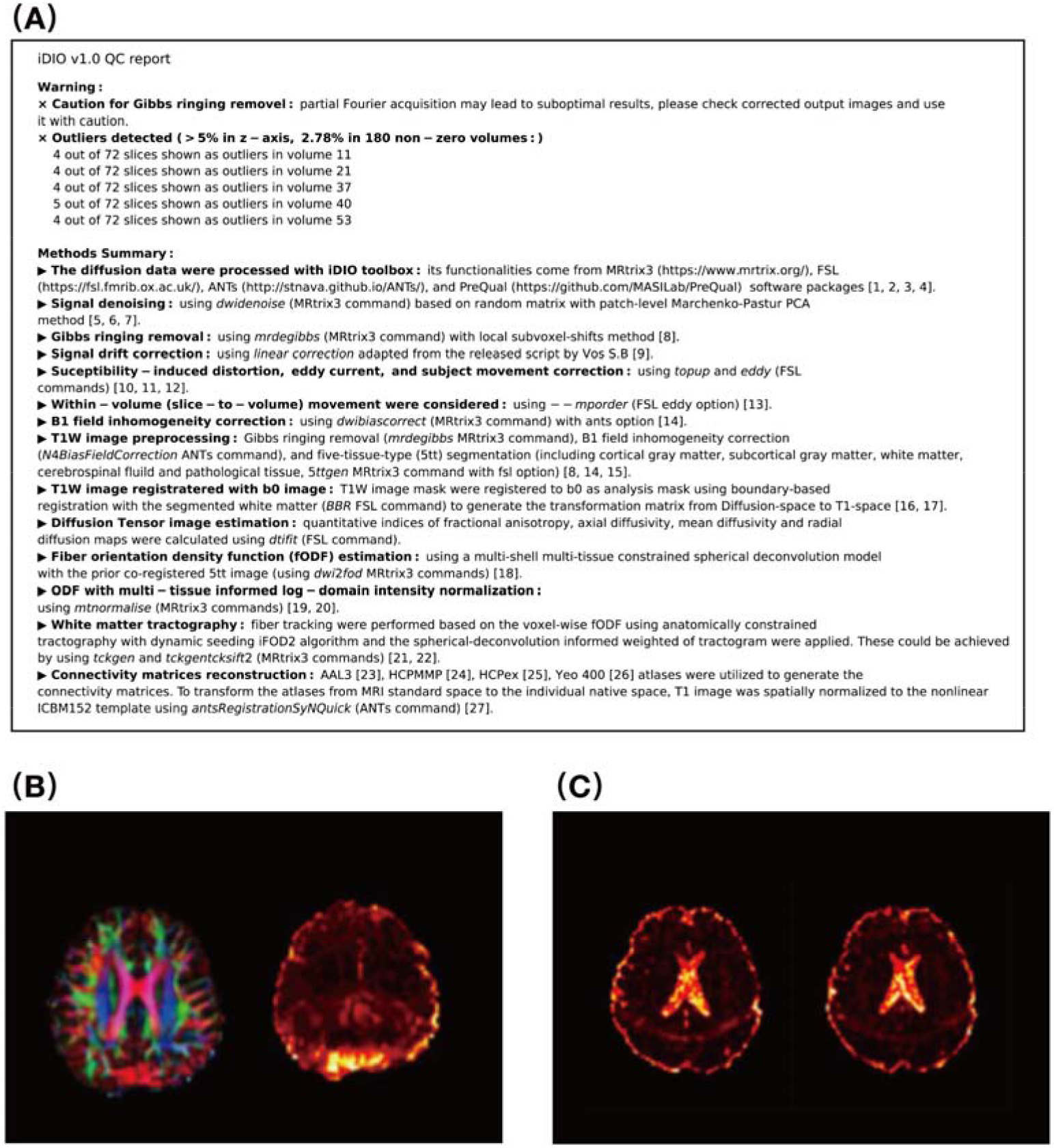
The summary report of iDIO, and the DEC and SSE map for visual check. (A) the report will remind the users for possible ill-usage in the warning part and an outlier summary for quick check. The DEC and SSE map is useful in identifying artefacts such as (B) vibration artefact and (C) fat chemical shifting.

### Test data

For demonstration, we applied iDIO on a dataset of 99 healthy participants scanned on the Siemens 3T MR scanner at National Yang Ming Chiao Tung University. High-resolution T1w images were acquired using a 3D Magnetization-Prepared Rapid Gradient Echo (MPRAGE) sequence (TR/TE = 2530/3.5 ms, TI = 1100 ms, FOV = 256 mm, flip angle = 7°, matrix size = 256 × 256, 192 sagittal slices, slice thickness = 1 mm, no gap). DWIs were acquired using simultaneous multiband accelerated spin-echo echo planar imaging (Setsompop et al., 2012; Auerbach et al., 2013) (TR/TE = 3235/109.2 ms, matrix size = 110 × 110, 72 contiguous slices, voxel size = 2 × 2 × 2 mm^3^, multiband factor = 3) with two b-values of 1000 s/mm^2^ (30 diffusion directions) and 3000 s/mm^2^ (60 diffusion directions), plus six interleaved b=0 images in two phase encoding directions (anterior to posterior and posterior to anterior). All studies were approved by the Institutional Review Board of the National Yang Ming University, Taipei, Taiwan.

We calculated the signal-to-noise ratio (SNR) of the DWIs following each step: S1) original input, S2) noise removal, S3) Gibbs ringing removal, S4) drifting correction, S5) susceptibility correction and eddy/motion correction and S6) bias field correction. The global SNR was estimated by the median of intra-brain voxel-wise SNR for each shell, which was defined as the mean divided by the standard deviation of the diffusion-weighted signal of the shell images. A repeated measure ANOVA was then conducted to compare among S1-S6.

## Results

### Data produced by iDIO

Table 1 lists all the output folders and files generated by iDIO, and the storage space required for the test data.

**Table 1.** Folders, files by iDIO. The storage cell is the size of files from a representative subject.

| Processing step | Folder name | Note | Storage occupied | Runtimes (min:s) |
| --- | --- | --- | --- | --- |
| BIDS_NIFTI | 0_BIDS_NIFTI | Rename files based on JSON file | 114.2 MB | <1s |
| DWI preprocessing | 1_DWIprep | Preparation files | 421 KB | <1s |
|  | 2_BiasCo | Denoise, Gibbs ringing removal, and drift correction | 2.5 GB | 7:40 |
|  | 3_EddyCo<sup>a</sup> | Susceptibility correction, eddy correction | 1.8 GB | 33:07 |
|  |  | bias field correction, and resize image |  |  |
| T1 preprocessing | 4_T1preproc | Gibbs ringing removal, bias field correction, brain extraction and registration to DWI space | 151 MB | 12:49 |
| DTI fitting | 5_DTIFIT | DTI fitting | 760.1 MB | 00:49 |
| CSD modeling | 6_CSDpreproc | CSD modeling | 1.6 GB | 01:16 |
| Network construction (including generating tracks* based on CSD FOD) | 7_NetworkProc | Generating tractogram, registration T1 to MNI to generated atlases in DWI space, weighted connectivity matrices | 8.2 GB | 81:34 |
| QC steps | 8_QC | QC report and stats file | 3.9 MB | 06:32 |
|  | Preprocessed_data | Preprocessed DWI and T1 images | 282.6 MB | <1s |
|  | Connectivity_Matrix | Atlas in T1 and DWI space, tractography assignment information, and connectivity matrices with length, mu weighted and SIFT weighted | 600 MB | <1s |
| Note: *including generating 10 million tracks based on CSD FOD.; The processing was performed using a local workstation with 128GB of memory, intel i9-7900X @3.30 GHz CPU with 20 cores and NVIDIA Quadro P4000 GPU. |  |  |  |  |

### Typical runtimes

Table 1 shows the runtime for each processing method considered. Step 3_EddyCo needed the longest preprocessing time (33 mins and 7 secs) with CUDA acceleration. The reconstruction of the brain network took the longest time (81 mins 34 secs), largely for generating the tractogram of 10 million streamlines.

### Data QC and SNR

The QC report from the complete iDIO preprocessing showed that two out of ninety-nine participants’ dMRI data were acquired without reverse phase data, and one was acquired with a problematic pair of phase encoding directions: one went anterior-posterior and the other went right-left, possibly caused by the manual entry error during the scan. Moreover, nine participants showed signal dropout in more than 3% of their DWI volumes. All the problematic data mentioned above were warned by iDIO’s log file and QC report; they are excluded from the SNR evaluation. Hence, in the end, eighty-seven participants’ image data were considered for the SNR evaluation.

Figure 4 shows the median values of the intra-mask SNR, signal, and noise, as computed for each subject at each processing stage. On the b=0 images, the group SNR increased from 9.56 ± 1.22 to 21.15 ± 1.73 after running the complete preprocessing (Figure 4A). Notably, the most significant SNR improvement occurred following 3_EddyCo. Such an increase in SNR mainly came from the significant reduction of the noise level, which was 14.13 ± 2.85 and 7.42 ± 1.05 before and after 3_EddyCo, respectively. On the DWIs, the SNR increase from the raw image to the denoise step was more prominent, especially at b=3000 (Figure 4C). Likewise, the SNR improvement was mainly driven by the reduced noise level after DWI denoising (from 8.68 ± 0.79 to 6.02 ±0.57). For both b-values, the repeated measures ANOVA revealed that the SNR, signal, and noise differed significantly between processing steps (p<0.001). Figure 5 shows the outcomes of the Post hoc test on the mean values, using a significant level of Bonferroni corrected p<0.001. Significant reductions of the signal were observed on the DWIs at both b-values after the bias correction step.

**Figure 4.**
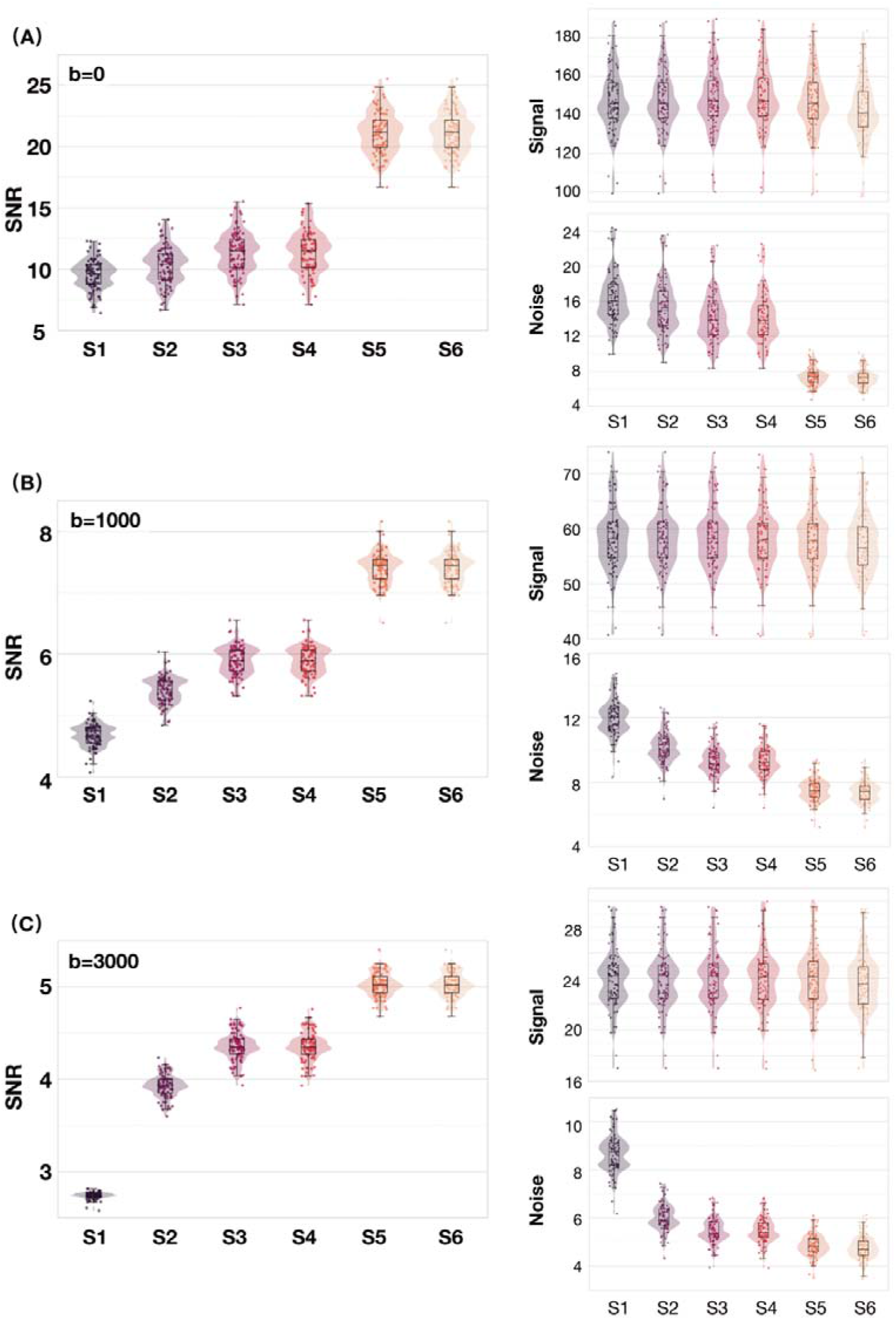
Median intra-brain SNR, signal, and noise of A) b=0, B) b=100, and C) b=3000 DWIs following per-processing step of iDIO. For a total of 87 participants with acceptable image quality (motion outlier were smaller than 3%), the standard iDIO preprocessing steps were applied in the order as follows: S1) Raw input images, S2) noise removal, S3) Gibbs ring removal, S4) signal drift correction, S5) correction for susceptibility-induced distortion, eddy current-induced distortion and subject movement, and S6) bias field correction. Each circle corresponds to the median value from a single subject. The global SNR was estimated by the median of intra-brain voxel-wise SNR for each shell, which was defined as the mean signal (Signal) divided by the standard deviation of the diffusion-weighted signal (Noise) of the images. For b0 and both b-values, the SNR, signal, and noise were differed significantly between processing steps (p<0.001).

**Figure 5.**
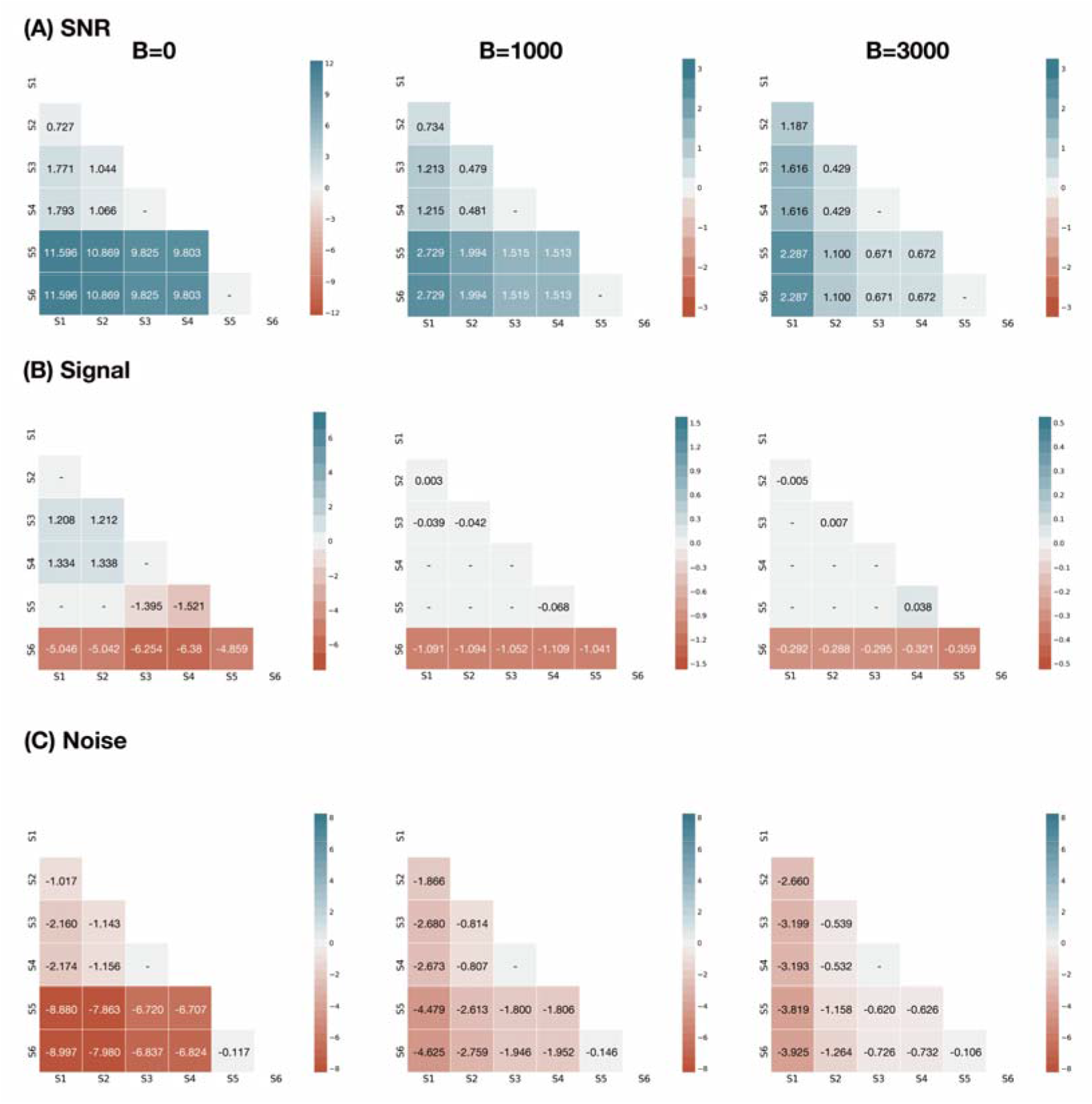
The mean differences of A) SNR, B) signal, and C) noise between each processing step. Significant differences in the post hoc test were shown in blue or red (p<0.001 Bonferroni corrected). Gray boxes with dash (-) symbols indicated the differences did not reach the significant level. S1) Raw input images, S2) noise removal, S3) Gibbs ring removal, S4) signal drift correction, S5) correction for susceptibility-induced distortion, eddy current-induced distortion and subject movement, and S6) bias field correction.

## Limitations

The present version of iDIO has several limitations. 1) iDIO does not have a Gaussian Process for diffusion-weighted signal modeling and outlier replacement (Andersson and Sotiropoulos, 2015). There is a need to implement a mechanism that could detect image artifacts such as signal loss, chemical shift, and motion in the preprocessed data. 2) iDIO is customized for shell-schemed dMRI data and hence does not support the sampling on a Cartesian grid, such as used in diffusion spectrum imaging (Wedeen et al., 2008). 3) iDIO recommends input data arranged in the BIDS format and information of image acquisition parameters in JSON. Data provided in ANALYZE or NIFTI without the corresponding JSON files might not be compatible due to the missing image header information. 4) Correction for susceptibility-induced distortion using field maps is not supported yet. Image without correction may lead to a misalignment with anatomical images, especially at the air-tissue boundary, which is detrimental to voxel-wise modeling and tractography. A potential solution is to take synthesized undistorted b=0 images as the reference images for distortion correction (Schilling et al., 2019), which will be included in the future release. 5) iDIO has a high demand for data storage space since it keeps nearly all the intermediate files for retrospective inspections on the data 6) Users need to install all the dependencies of iDIO manually, which is still not handy for non-experts. A potential solution is to distribute everything as a whole using containers, such as Docker or Singularity, at the price of large container size. 7) iDIO will be developed continuously to be more comprehensive; new features such as reduction of Rician noise and correction for gradient nonlinearity (Tax et al., 2021) are needed.

In summary, iDIO provides a semi-automated way for preprocessing dMRI data. We minimize the required inputs to prevent manual errors and record the processing arguments in the log files. Both functions facilitate the applications of data preprocessing, quality control, and management. The source codes and documentation of iDIO are publicly available at https://github.com/iDIO4dMRI/iDIO.

## Acknowledgements

The authors would like to acknowledge Dr Chun-Hung Yeh for the discussions on pipeline development and manuscript editing. Professor C-P. Lin was supported by the Ministry of Science and Technology (MOST) of Taiwan (grant number: MOST 111-2321-B-A49-003, MOST 111AT38A), and Veterans General Hospitals and University System of Taiwan Joint Research Program (Grant number VGHUST110-G1-3-3).

## Notes

https://github.com/iDIO4dMRI

### Competing Interest Statement

The authors have declared no competing interest.

